# Named Entity Recognition of Pharmacokinetic parameters in the scientific literature

**DOI:** 10.1101/2024.02.12.580001

**Authors:** Ferran Gonzalez Hernandez, Quang Nguyen, Victoria C. Smith, José Antonio Cordero, Maria Rosa Ballester, Màrius Duran, Albert Solé, Palang Chotsiri, Thanaporn Wattanakul, Gill Mundin, Watjana Lilaonitkul, Joseph F. Standing, Frank Kloprogge

## Abstract

The development of accurate predictions for a new drug’s absorption, distribution, metabolism, and excretion profiles in the early stages of drug development is crucial due to high candidate failure rates. The absence of comprehensive, standardised, and updated pharmacokinetic (PK) repositories limits pre-clinical predictions and often requires searching through the scientific literature for PK parameter estimates from similar compounds. While text mining offers promising advancements in automatic PK parameter extraction, accurate Named Entity Recognition (NER) of PK terms remains a bottleneck due to limited resources. This work addresses this gap by introducing novel corpora and language models specifically designed for effective NER of PK parameters. Leveraging active learning approaches, we developed an annotated corpus containing over 4,000 entity mentions found across the PK literature on PubMed. To identify the most effective model for PK NER, we fine-tuned and evaluated different NER architectures on our corpus. Fine-tuning BioBERT exhibited the best results, achieving a strict *F*_1_ score of 90.37% in recognising PK parameter mentions, significantly outperforming heuristic approaches and models trained on existing corpora. To accelerate the development of end-to-end PK information extraction pipelines and improve pre-clinical PK predictions, the PK NER models and the labelled corpus were released open source at https://github.com/PKPDAI/PKNER.

## 1 Introduction

Bringing a new chemical compound to the market is an extremely costly process, which has been estimated between $161m and $4.5bn [Schlander et al., 2021]. Meanwhile, over 90% of drug candidates fail after entering phase I clinical trials [Wong et al., 2019, DiMasi et al., 2016]. Accurate predictions of candidate drug properties at an early stage are critical for improving the efficiency of this process. To elicit the desired effect, candidate drugs must reach a specific concentration at the target site of the body over a certain time period [Morgan et al., 2012]. Predicting whether candidate drugs will reach the desired concentration over a certain period at the target site requires understanding the processes of absorption, distribution, metabolism and excretion (ADME) of drugs from the human body.

Pharmacokinetic (PK) parameters quantify the ADME processes of chemical compounds through numerical estimates. Accurate estimation of drugs’ PK parameters is crucial for drug development research [Morgan et al., 2012]. Mechanistic models have been widely used to predict the PK parameters of candidate drugs before they are tested in humans. However, a significant proportion of those candidates still fail due to PK complications found during the clinical phases [Palmer, 2003]. Hence, improving PK predictions of candidate compounds before they are given to humans is crucial for assessing candidate prospects and optimising the drug development pipeline.

One of the main challenges in improving PK predictions for chemical compounds is the lack of comprehensive and standardised PK repositories [Grzegorzewski et al., 2021, Gonzalez Hernandez et al., 2021]. Although existing open-access databases collect information ranging from chemical structure to a summary of PK publications, they typically only report sparse PK information explicitly [Grzegorzewski et al., 2021, Wong et al., 2019, Gonzalez Hernandez, 2022]. Consequently, researchers must search and curate PK estimates from scientific literature before pre-clinical predictions can be made [Grzegorzewski et al., 2021, Lombardo et al., 2018]. The vast and continually increasing number of PK publications, coupled with the extensive amount of PK information locked in scientific articles, limits our ability to efficiently find and curate comprehensive datasets manually [Wong et al., 2019]. Thus, despite the potential PK data stored in scientific articles, efficiently exploiting this resource remains a significant challenge in drug development.

Automated text mining approaches can aid researchers in extracting information from the scientific literature more efficiently. Recognising entities of interest is a crucial step in information extraction pipelines that enables subsequent downstream tasks such as relation extraction or entity linking. In this study, we focus on the initial step towards automated extraction of PK parameter estimates from the scientific literature, Named Entity Recognition (NER). Developing systems that can identify mentions of PK parameters in scientific text is crucial for end-to-end PK extraction as well as characterising drug-drug interactions (DDIs), as many interactions are reported by mentioning their effect on specific PK parameters [Kolchinsky et al., 2015]. However, PK NER remains a challenging task since there are multiple PK parameter types and their mentions are often highly variable across the scientific literature, involving the frequent use of acronyms and long textual spans [Wu et al., 2013]. Additionally, the scarcity of annotated resources limits the development of effective NER models that can deal with this diversity. In this work, we tackle these challenges by developing annotated corpora and machine-learning models for effective PK NER.

## 2 Methodology

### 2.1 Corpus construction

A protocol was established to generate corpora of labelled sentences that allowed training and evaluation of PK NER models. The final corpus is referred to as the PK-NER-Corpus and can be found at https://zenodo.org/records/4646970 [Gonzalez Hernandez, 2024].

#### Source

To create a candidate pool for sentence annotation, the pipeline described in Figure 1 was applied. A PubMed search for *“pharmacokinetics”* was initially conducted to retrieve articles. Using the pipeline from Gonzalez Hernandez et al. [2021], 114,921 relevant publications reporting PK parameters were identified. Out of these, 10,132 articles (8.82%) were accessible in full text from the PMC OA subset^1^, while only abstracts were available for the rest. Both, abstracts and full-text articles were downloaded in XML format from PubMed^2^ and PMC^3^ FTP sites. The PubMed Parser [Titipat and Acuna, 2015] was used to parse the XML files, and paragraphs from the introduction section were excluded. The scispaCy sentence segmentation algorithm [Neumann et al., 2019] split abstracts and paragraphs into sentences. The resulting sets were the abstract pool with over a million sentences and the full-text pool with 721,522 sentences. To create a balanced candidate pool for ML model training and evaluation, 721,522 instances were randomly sampled from the abstract pool and combined with full-text sentences, resulting in a balanced pool of 1,443,044 sentences, referred to as the candidate pool. All labelled sentences in the corpus construction were sampled from the candidate pool.

**Figure 1:**
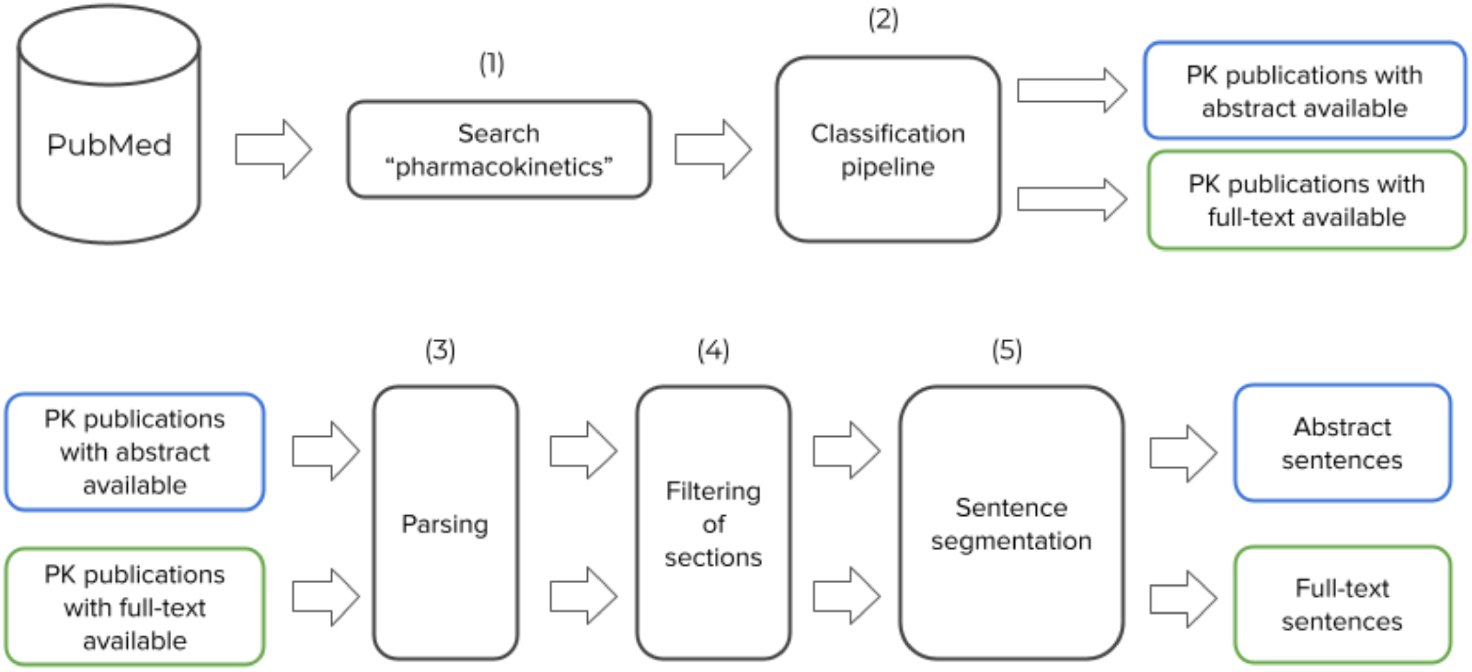
Flow diagram showing the main processes involved to generate a pool of candidate sentences for NER labelling. (1) Search for ‘pharmacokinetics’ in PubMed and (2) run binary classification pipeline to filter abstracts containing PK parameters. (3) Parse XML abstract and full-text documents, and (4) filter out introduction sections. Finally, (5) segment each paragraph into sentences to generate the final corpus of PK sentences.

#### Annotation

The team responsible for the annotation involved twelve annotators with extensive PK expertise and familiarity with the different parameters and study types in the PK literature. To ensure consistency in the annotation process, each annotator initially labelled a small set of 200 examples to identify sources of disagreement. The team then discussed which parameters to include and how to define span boundaries using the PK ontology from Wu et al. [2013] as a reference. Annotation guidelines were provided to annotators before they began the labelling task, and were updated as new challenging examples were resolved during the annotation process. Details about the annotation interface and guidelines can be found in Appendix A. Training, development and test sets were developed to train and evaluate different NER pipelines.

#### Training set

Training effective NER models for new entity types often requires a large number of annotated samples with diverse spans to account for the variability of surface forms and contexts of use [Wang et al., 2018]. However, the sampling strategy followed by the test and development sets resulted in a low proportion of sentences containing PK entities (16.4%). To generate an effective training dataset, two main approaches were sequentially applied to selectively sample informative sentences for PK NER while reducing annotation efforts:

##### 1. Heuristic labelling

The rule-based model described in Appendix B was applied to all the sentences within the candidate pool. From those sentences that contained matches from the rule-based model, a total of 300 were randomly chosen to form an initial training set with a substantial number of entity mentions. To enhance the quality of this set, the annotators corrected the labels generated by the rule-based model. Following correction, 86.67% of the sentences retained PK mentions, although adjustments were often needed for their span boundaries. Subsequently, an initial scispaCy NER model (*en_core_sci_md* [Neumann et al., 2019]) was trained on this dataset.

##### 2. Active learning

After training the initial scispaCy model, it was used to identify spans from the candidate pool that were most informative for model training. Utilising the active learning interface from Prodigy [ExplosionAI, 2021], which presents candidate spans to annotators based on model uncertainty, annotators provided binary labels denoting the correctness of suggested spans. During the active learning process, the model underwent updates in a loop after every set of 10 annotated sentences. After obtaining binary labels, a final round of annotation was conducted to label any additional spans present in the sentences and correct span boundaries. Following this protocol, a total of 2800 sentences with a large number of PK entity mentions were labelled. Further details on the Active Learning protocol can be found in Appendix C.

#### Test and development sets

The development and test sets were generated by randomly sampling sentences from the candidate pool without replacement, to preserve the distribution of sentences found in PK articles. In total, 1,500 and 500 sentences were selected for the test and development sets, respectively. Then, each sentence in the development and test sets followed a two-stage procedure of (1) initial annotation by one expert and (2) review and standardisation of span boundaries by at least two additional experts (similar to [Hope et al., 2020]). This process was carried out in batches of 200 sentences. After each batch, sources of disagreement were discussed, and annotation guidelines were updated.

#### Inter-annotator agreement (IAA)

We selected pair-wise F1 as the main metric for measuring IAA in NER [Hripcsak and Rothschild, 2005, Deleger et al., 2012]. IAA was computed for each pair of annotators and F1 was obtained by treating the labels of one annotator as ground truth and the other as the system prediction. All annotators independently labelled a total of 200 sentences from the test set, used to derive the IAA. This exercise was done with the last batch of the test set when guidelines had already been updated multiple times, but no corrections were performed before computing the IAA.

#### External dataset validation

We utilised the PK Ontology and its corresponding corpus developed by Wu et al. [2013] for external validation. This corpus, referred to as PK-Ontology-Corpus, comprises 541 abstracts manually labelled, encompassing the annotation of key terms, sentences related to Drug-Drug Interactions (DDI), and annotated DDI pairs. The abstracts originated from four study types: clinical PK, clinical pharmacogenetics, *in vivo* DDI, and *in vitro* DDI studies. One of the annotated key terms in the PK-Ontology-Corpus was PK parameters. The NER models developed in this study were also evaluated in the PK-Ontology-Corpus, which allowed for assessing model performance in different study types, including several DDI sentences and detecting differences in the annotation criteria.

### 2.2 Models

#### Rule-based system

Given the PK expertise of the annotation team, a set of rules was generated to develop a rule-based model covering well-known PK parameters and their primary surface forms and acronyms. The model was implemented using the entity ruler from spaCy, which requires a set of token-level patterns and can incorporate rules regarding part-of-speech (POS) and dependency labels. ScispaCy [Neumann et al., 2019] was used as a base tokeniser, POS tagger and dependency parser to incorporate the token-level patterns into the model. Developing the list of terms and rules was an iterative process performed together with the annotation team, and rules were updated assessing their performance on the development set. For a list of patterns see Appendix B.

#### BERT

The Transformer architecture has emerged as state-of-the-art for NLP tasks [Vaswani et al., 2017]. In this study, pre-trained BERT models were fine-tuned to perform PK NER [Devlin et al., 2018]. We added a task-specific layer (fully-connected + softmax) to map output token embeddings from BERT models to BIO labels [Campos et al., 2012]. Two pre-trained models were compared: *BERT*_*BASE*_ [Devlin et al., 2018] which was pre-trained on general-domain English text, and BioBERT v1.1 [Lee et al., 2020] which was further pre-trained on PubMed articles. Models were implemented in PyTorch [Paszke et al., 2017] using the Transformers library [Wolf et al., 2019].

BERT tokenizers split each input sentence into sub-word tokens, each associated with a BIO label. The model was trained to minimise categorical cross-entropy loss. Both BERT and classification layer parameters were fine-tuned during 20 epochs. The model’s performance was evaluated on the development set at the end of each epoch, saving the state with the highest entity-level F1 score. We used the Adam optimizer with a linear weight decay of 0.05 and a dropout probability of 0.1 on all layers. We used a batch size of 16 and the learning rate was grid-searched, with *µ* = 3*e*^*−*5^ yielding the best performance. The maximum sequence length was set to 256 to cover most training instances. During inference, sentences with over 256 tokens were split, and predictions were re-joined after BIO label assignments. Experiments ran on a single NVIDIA Titan RTX (24GB) GPU.

#### ScispaCy

The scispaCy model was also fine-tuned to perform NER of PK parameters. ScispaCy is built on top of spaCy but focuses on biomedical and scientific text processing [Neumann et al., 2019]. In this work, all components from the scispaCy pipeline (*en_core_sci_md*) were reused, and the NER layer was trained from scratch. Analogous to the BERT pipelines, models were trained for 20 epochs and the state of the model with the best performance on the development set was saved. The rest of the hyperparameters were kept identical to Neumann et al. [2019].

#### Evaluation

We computed precision and recall, and derived F1 score for comparing model performance. To determine true positives we used both, strict and partial matching. Strict matching requires complete overlap in entity boundaries between predictions and annotations while partial matching considers instances where system predictions partially overlap with annotated entities. Both strict and partial matching metrics were computed using the *nervaluate* library^4^.

## 3 Results and Discussion

### 3.1 Corpus statistics

The main statistics for the PK-NER-Corpus are shown in Table 1. Since the evaluation sets randomly sampled sentences from PK articles, the proportion of sentences containing PK parameter mentions was only 16.40%. Despite preserving the distribution of sentences in which PK NER algorithms might be applied, fewer entity mentions were present in the evaluation sets. On the other hand, 64.25% of sentences in the training set contained mentions of PK parameters, resulting in many entity mentions. This difference in the distribution of parameter mentions was due to the active learning sampling protocol selecting sentences with a higher proportion of entity mentions. Additionally, while we randomly sampled sentences from the abstract or full-text section in the evaluation sets, the active learning protocol selected a higher proportion of sentences from the full text (79.56%).

**Table 1:** Corpus statistics of the PK-NER-Corpus stratified by the training, development and test sets The statistics of our external evaluation corpus (PK-Ontology-Corpus) from Wu et al. [2013] are shown in Table 2.

| Dataset | Sentences | Entity mentions | Sentences with PK mentions (%) | Full-text sentences (%) |
| --- | --- | --- | --- | --- |
| Training | 2800 | 3680 | 64.25 | 79.46 |
| Development | 500 | 149 | 16.40 | 50.8 |
| Test | 1500 | 390 | 16.40 | 50.8 |

**Table 2:** Corpus statistics of the PK-Ontology-Corpus stratified by the training and test sets.

| Dataset | Sentences | Entity mentions | Sentences with PK mentions (%) |
| --- | --- | --- | --- |
| Training | 4008 | 1478 | 23.68 |
| Test | 1021 | 377 | 25.27 |

### 3.2 Effects of Active Learning

To evaluate the effects of active learning we performed the following experiment. The development set (*n* = 500) was used as an example of an annotated set randomly sampled while 500 sentences from the training set collected with active learning were randomly sampled to perform a fair comparison. Ten separate runs with different random seeds were performed. The active learning experiment randomly sampled a different subset of sentences from the training set and randomly initialised the classification layer parameters in each run. The BioBERT model was trained for five epochs with a learning rate of 3*e*^*−*5^, and the final model was applied to the test set at the end of each run.

Figure 2 show the results of these experiments. Training the BioBERT model with the active learning dataset resulted in over 7% increase in the median F1 score for strict matching compared to training with randomly sampled sentences. These results suggest that the protocol used to generate the training set highly benefited the model performance compared to randomly sampling sentences. Most of this benefit is the consequence of an improved recall, suggesting that the active learning dataset contains a wide variety of PK spans not covered by the random sampling dataset. Considering the frequency of named entities in each dataset (i.e. only 16.4% of sentences mentioned PK parameters in the randomly sampled datasets), it is likely that the selective sampling approach implemented for this task was particularly beneficial for covering a wider variety of relevant spans.

**Figure 2:**
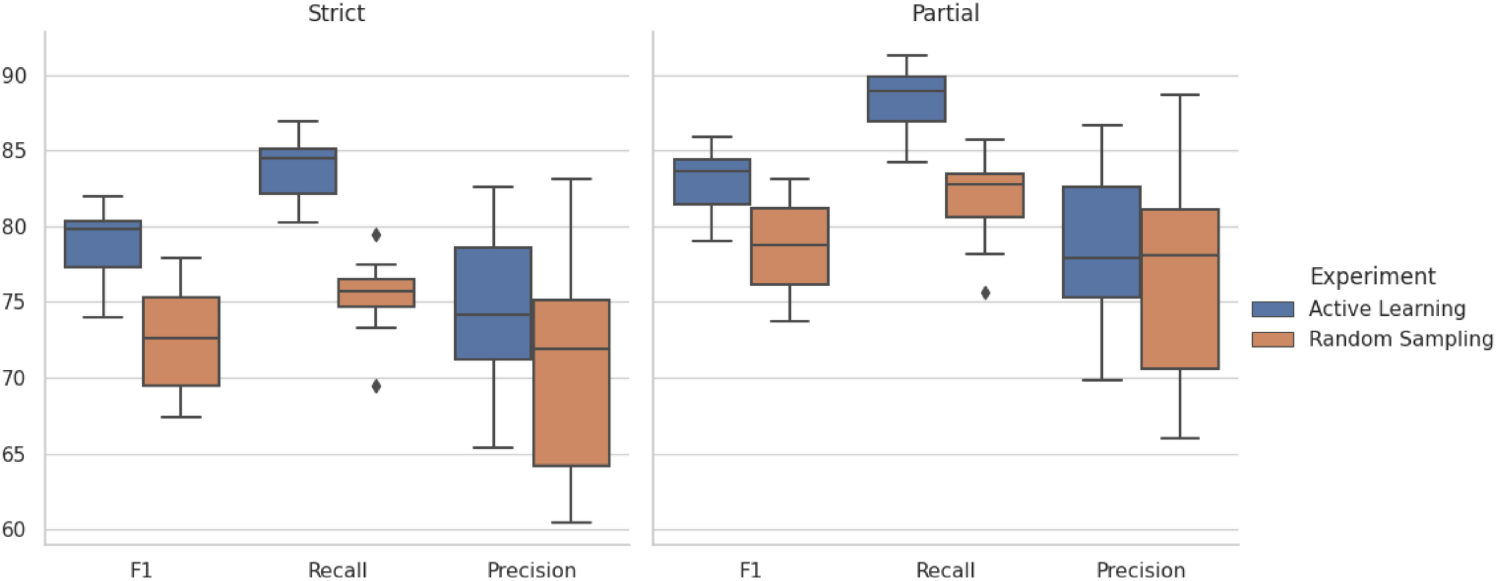
Distribution of F1, Recall and Precision scores for the *Active Learning* and *Random Sampling* datasets (n=500 sentences) after 10 runs with different random seeds. The left and right panels display the scores considering strict and partial matching of entities, respectively.

### 3.3 Model performance

Table 3 summarises the main results on the test set. The results showed that the rule-based model could not efficiently cover the diversity of PK parameter mentions annotated by field experts on the test set, with a strict F1 score below 50%. Some of the main challenges of the rule-based approach were (1) the great variety of PK parameter types, which limited the pipeline’s recall, (2) the presence of complementary terms within PK spans that were difficult to encapsulate with rules (e.g. total body clearance), (3) acronyms highly dependent on context (e.g. “F” for bioavailability, “AUC” for the area under the concentration-time curve). Notably, there was a large difference in precision between strict and partial matches (over 15%). This is a consequence of challenge (2), where rules often detected the primary PK term, but complementary terms determining the parameter sub-type were missed.

**Table 3:** Results on the test set for different NER models. Metrics are reported at the entity level using strict and partial matches.

| Model | Strict |  |  | Partial |  |  |
| --- | --- | --- | --- | --- | --- | --- |
| | P | R | $F_1$ | P | R | $F_1$ |
| Rule-based | 52.8 | 43.59 | 47.75 | 69.25 | 57.18 | 62.64 |
| ScispaCy | 77.09 | 82.82 | 79.85 | 80.91 | 86.92 | 83.81 |
| BERT | 81.47 | 87.72 | 84.48 | 84.92 | 91.43 | 88.05 |
| BioBERT | **90.49** | **90.26** | **90.37** | **92.54** | **92.31** | **92.43** |

The machine learning pipelines significantly outperformed the heuristic model with over 30% gain on the strict F1 score and particularly large improvements on recall. Furthermore, as it has been previously reported [Weber et al., 2021], it was observed that the models based on BERT provided substantial performance benefits in comparison to the scispaCy model. From the test set predictions, it was observed that the scispaCy pipeline was x10 faster at inference time on CPU in comparison to running BERT models on a single GPU. Therefore, we also released the fine-tuned scispaCy pipelines open-source ^5^.

The BioBERT model outperformed the BERT model pre-trained on general-domain English text, especially on strict entity matching. Specifically, BioBERT provided a large gain (+ 9%) on the pipeline precision in comparison to all the other models. This result suggests that domain-specific pre-training is crucial for effective PK NER.

### 3.4 Performance on External Corpus

The BioBERT model fine-tuned on the PK-NER-Corpus was applied to the test set of the PK-Ontology-Corpus, without any further training on this dataset. Surprisingly, the model reported a competitive strict F1 score of 74.52% without any training on that dataset (See Table 4). We observed a substantial increase from strict to partial matching (81.10%), which indicates that the main PK terms are often captured, but disagreements on span boundaries limit the model performance on this dataset. This difference might be explained by discrepancies in the span boundaries annotated between the PK-NER-Corpus and the PK-Ontology-Corpus.

**Table 4:** Results obtained on the external PK-Ontology-Corpus test set after training BioBERT on the PK-NER-Corpus.

| Training corpus | Strict |  |  | Partial |  |  |
| --- | --- | --- | --- | --- | --- | --- |
| | P | R | $F_1$ | P | R | $F_1$ |
| PK-NER-Corpus | 77.05 | 72.15 | 74.52 | 83.85 | 78.51 | 81.10 |

<sup>5</sup> https://github.com/PKPDAI/PKNER

Although different research groups developed both datasets, it was observed that the models trained on PK-NER-Corpus transferred well to the annotation criteria used in Wu et al. [2013]. However, when we trained a BioBERT model with the PK-Ontology-Corpus and applied it to the test set of the PK-NER-Corpus, the resulting strict matching F1 was 66.13%. Since the PK-Ontology-Corpus [Wu et al., 2013] used abstracts deliberately chosen (e.g. midazolam PK abstracts), models trained on this corpus probably learned some features specific to these types of articles but not transferable to PK studies from other types of compounds.

## 4 Conclusion and future work

This work presented a new corpus to train and evaluate NER models to detect mentions of PK parameters in the scientific literature. A variety of models were compared, and fine-tuning BioBERT resulted in the best performance on PK NER with over 90% F1 score on strict entity matching. Domain-specific pre-training with transformers was crucial to obtain optimal performance. Machine learning models largely outperformed the rule-based model, potentially due to the high diversity in PK parameter surface forms and the strong importance of context tokens to determine PK entities.

The active learning protocol helped accelerate the curation of PK data while improving the information provided by labelled sentences compared to random sampling. A variety of approaches have been applied for active learning in NER [Shen et al., 2017, 2004, Siddhant and Lipton, 2018]. For instance, Bayesian approaches have recently shown promising results [Siddhant and Lipton, 2018], although their application comes with computation costs. It is still unclear which active learning approaches are most beneficial to make efficient use of a model in the loop. In this study, many approaches are left for exploration. For instance, using transformer-based models in the loop instead of scispaCy, using diversity sampling or applying other criteria to estimate model uncertainty. However, the framework developed with Prodigy allowed for fast annotations that reduced the labelling load, and the samples selected for annotations provided diverse and challenging spans that resulted in larger information gains than samples randomly sampled.

Finally, the best-performing model showed good generalisation to various study types when applied to external annotated corpora and validated its potential application to improve the characterisation of DDIs. The experiment results indicate that NER models trained on the PK-NER-Corpus generalise better to unseen PK publications than those trained on existing corpora. Overall, we believe that these resources can become crucial in developing end-to-end PK information extraction pipelines, improving the characterisation of drug-drug interactions, and ultimately helping to improve PK pre-clinical predictions. To this end, we release our models and annotated corpora at https://github.com/PKPDAI/PKNER.

### Appendices

#### A Annotation

An interface was developed to annotate PK entities at the sentence level using the commercial tool Prodigy [ExplosionAI, 2021]. Figure 3 shows the main components of the interface. Each annotation instance was displayed in a single sentence, and field experts were asked to highlight those spans of text relating to PK parameters. If annotators were unsure about how to annotate a specific instance, they could flag the example for review. After annotations were performed, each sample stored the character-level boundaries and entity type for each annotated span (e.g. entity: “PK”, start character: 15, end character: 20).

**Figure 3:**
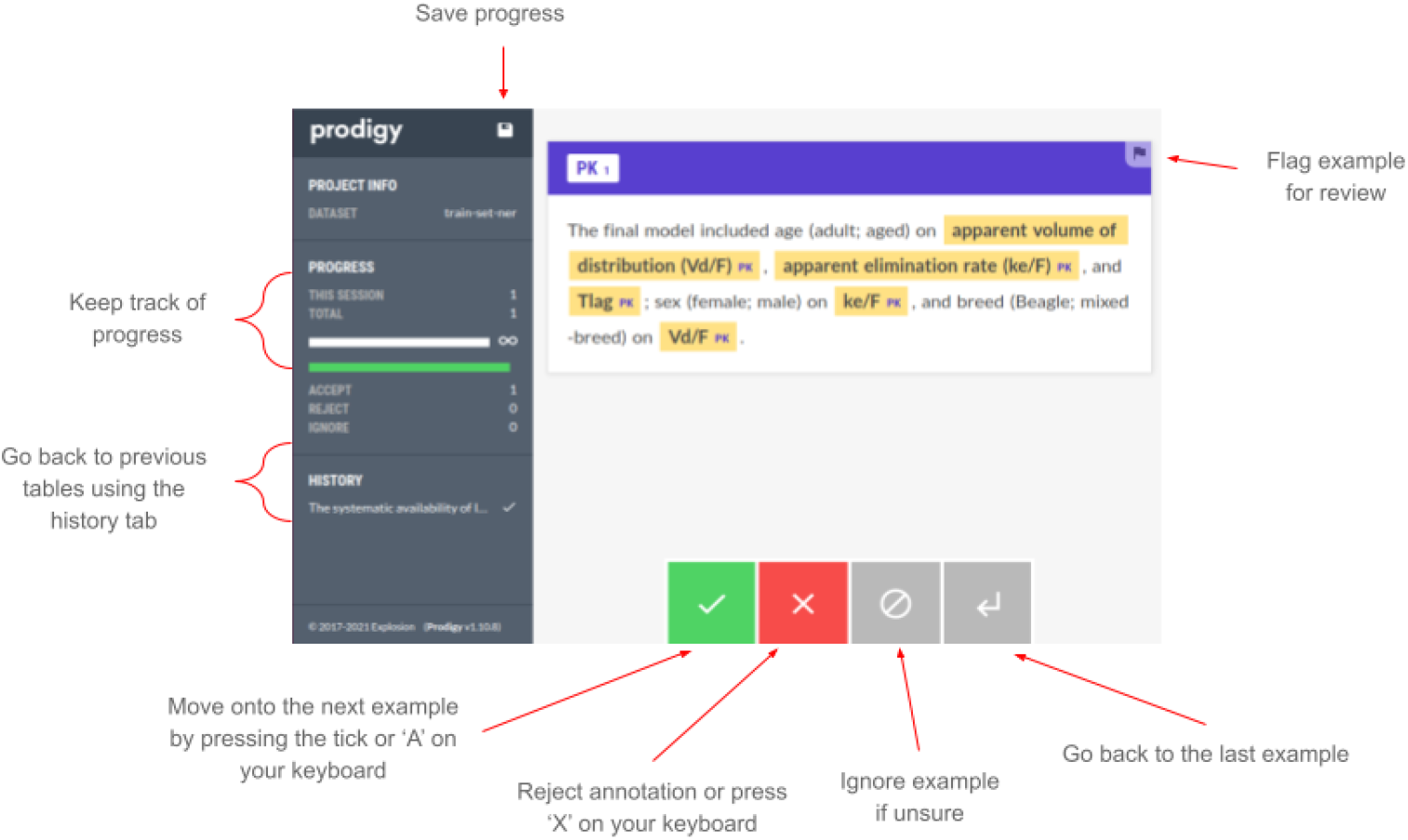
Screenshot of the interface used to annotate PK entities from scientific text. The example displays a single sentence after being annotated.

Explicit and detailed guidelines were required to ensure consistency in various decisions, especially those regarding span boundaries. The final annotation guidelines can be found at https://github.com/PKPDAI/PKNER/blob/main/guidelines/guidelinesner.pdf.

Amongst many others, some common doubts arising when annotating PK spans included:

- Which noun modifiers should be included as part of the PK span: *renal clearance* or *renal clearance*; *mean clearance* or *mean clearance*.
- How to label abbreviated forms: *oral clearance (CL/F)* or *oral clearance (CL/F), volume of distribution (V*_*d*_) or *volume of distribution (V*_*d*_).
- Ratios of PK and PD parameters: *AUC/MIC* or *AUC/MIC*.
- Whether to label parameter mentions appearing inside equations.

The resolution of which parameters to include and how to define span boundaries was discussed with the team, and decisions were based on the expected applications of the models developed, i.e. numerical PK parameter extraction and characterisation of drug-drug interactions (DDIs). Annotation guidelines were provided to annotators before starting the labelling task. As new challenging examples appeared and conflicting instances were resolved, guidelines were updated accordingly.

#### B NER Patterns

Below is the final list of spaCy patterns^6^ included to recognise PK parameters through rules. If more than one pattern generated overlapping spans, the longest span was selected.

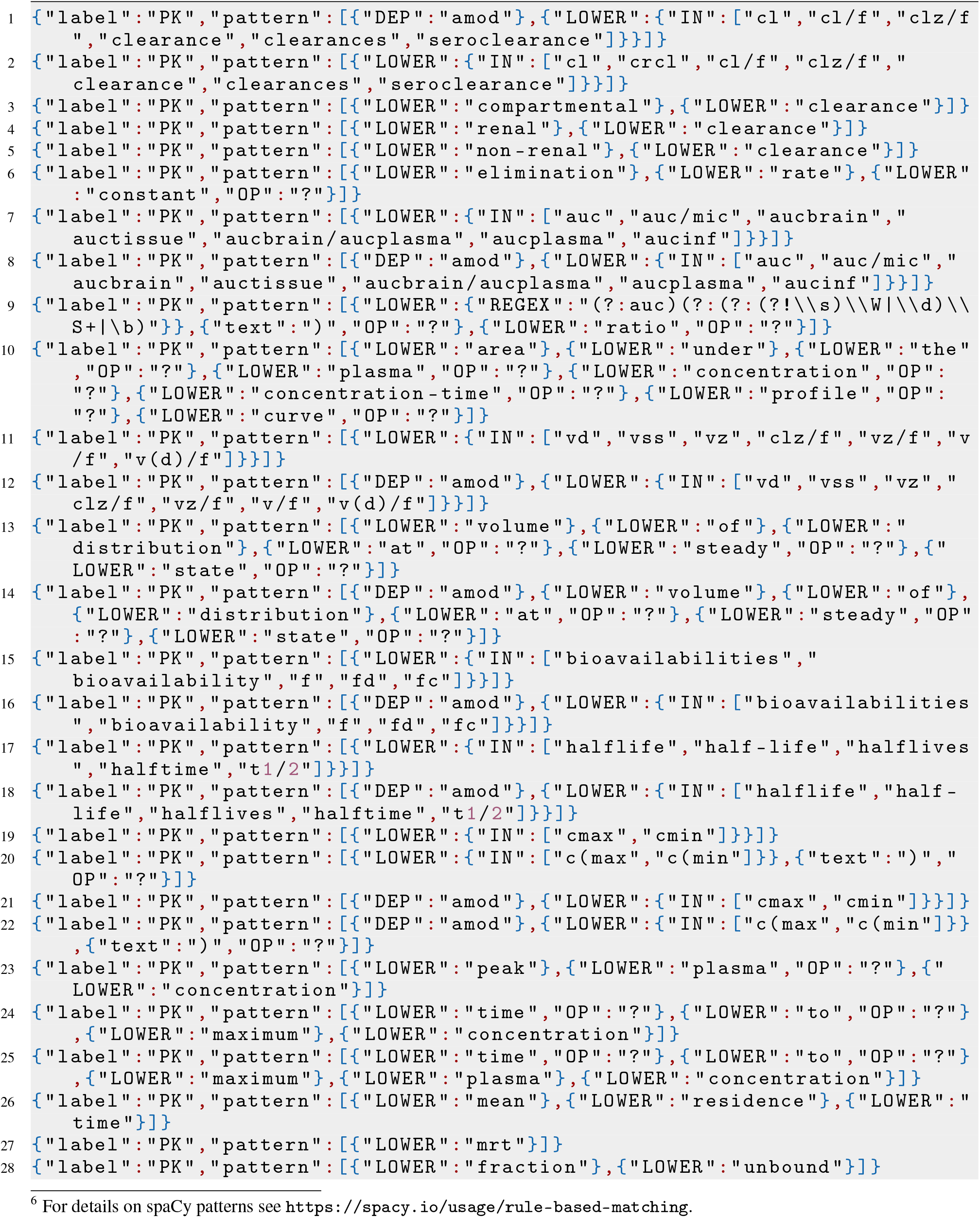
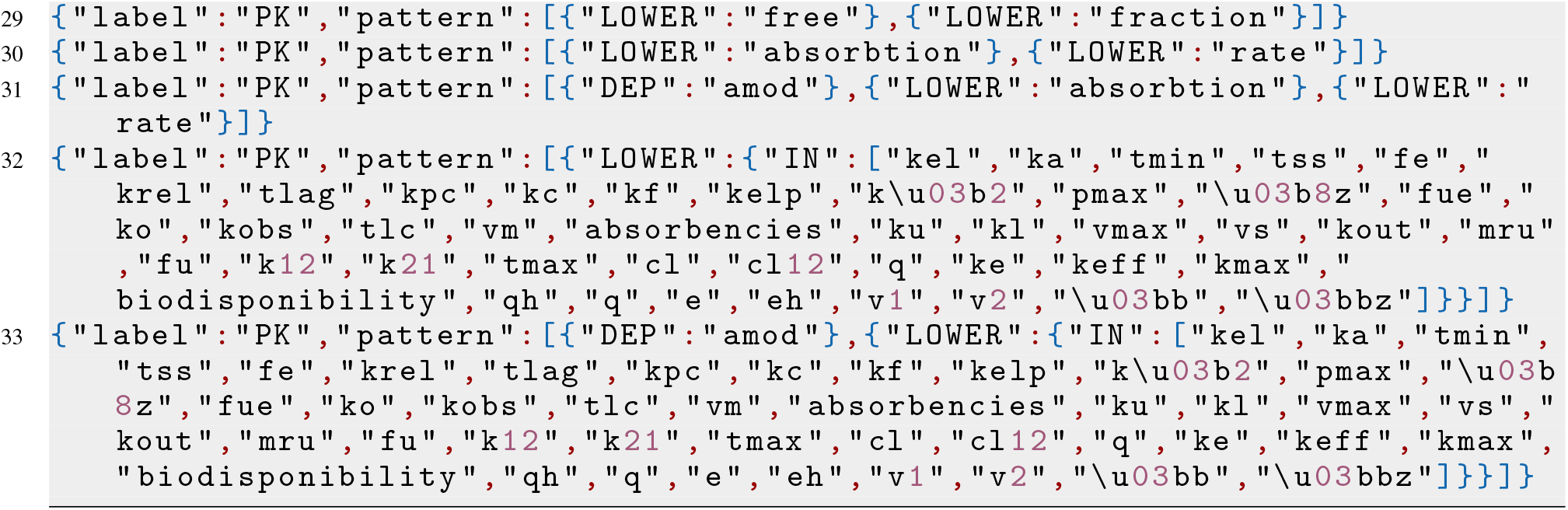

#### C Active Learning Protocol

As shown in Figure 4 (A), the scispaCy model (en_core_sci_md Neumann et al. [2019]) was trained with the initial dataset of 300 sentences (see section 4.2.1.4 for details on the model and training approach). After training the scispaCy pipeline on the initial dataset, the Prodigy ner.teach framework was used to suggest those spans where the model was most uncertain about based on the token-level softmax scores given by the NER model (B and C Figure 4). Specifically, Prodigy uses beam search to select the most uncertain spans over the sequence of token-level probabilities. The default settings from the Prodigy *ner*.*teach* framework were used (e.g. update approach and frequency). Annotators were asked to accept or reject spans predicted by the model, and based on their answers; the model was updated in the loop after every batch of ten binary annotations (D and E Figure 4).

**Figure 4:**
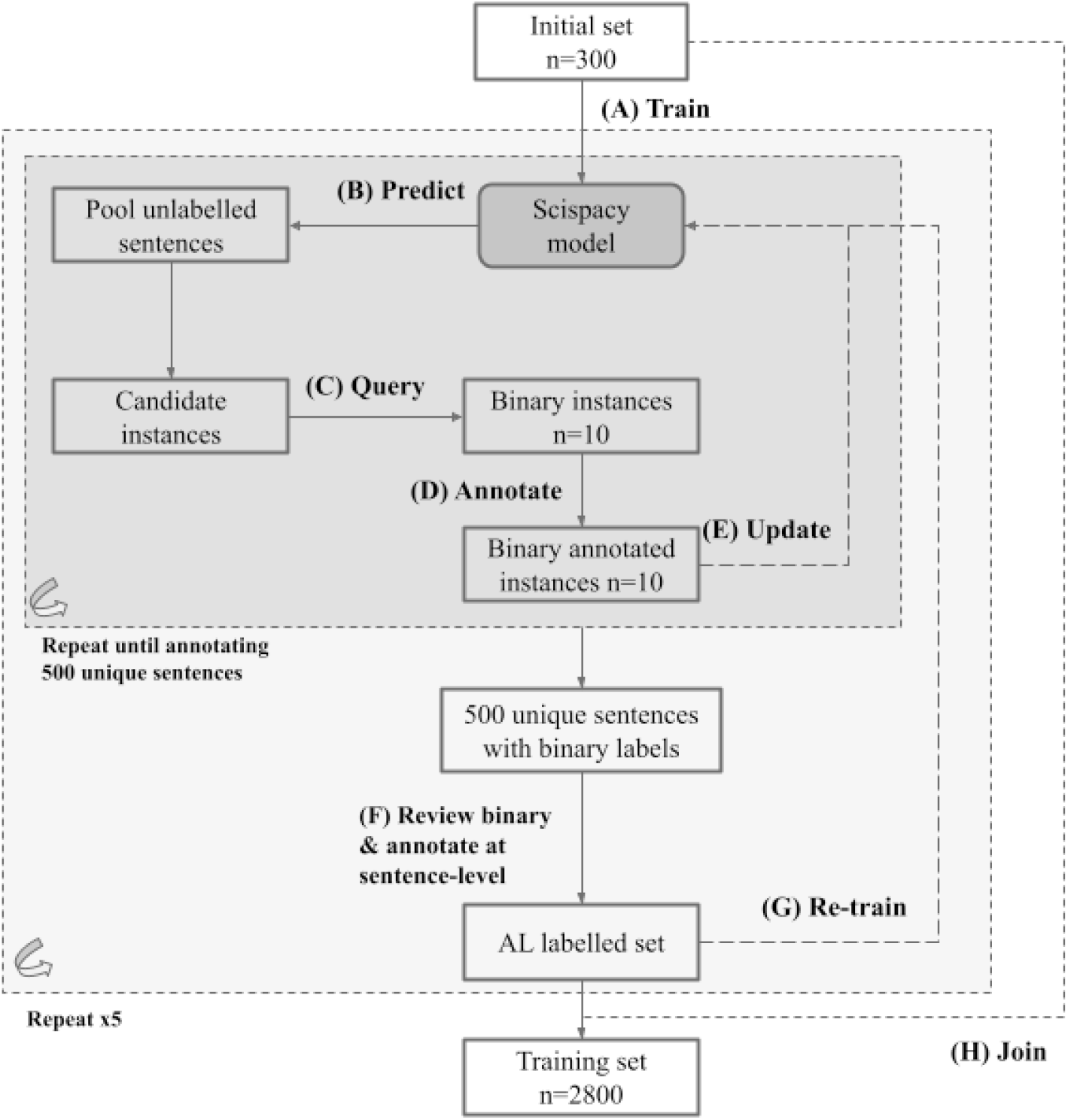
Schematic representation of the approach used to label instances for the training set by using and updating scispaCy NER model in the loop.

Instead of using large language models like BERT, the scispaCy pipeline allowed fast iterations and updates with the model in the loop, which made it preferable for AL purposes. This procedure was performed for 2500 sentences by a single annotator. Once binary annotations had been performed, another annotator reviewed each of the sentences with binary annotations and highlighted other spans if present in the sentence (step F in Figure 4). The model used for AL was re-trained (not updated) from scratch after every batch of 500 sentences was annotated (Figure 4, step G). Finally, the dataset collected with AL approaches was joined with the initial rule-based dataset, generating a training set of 2800 sentences (step H Figure 4).

## Footnotes

1 https://www.ncbi.nlm.nih.gov/pmc/tools/openftlist/

2 https://www.nlm.nih.gov/databases/download/pubmed_medline.html

3 https://ftp.ncbi.nlm.nih.gov/pub/pmc/

4 https://github.com/MantisAI/nervaluate

5 https://github.com/PKPDAI/PKNER

6 For details on spaCy patterns see https://spacy.io/usage/rule-based-matching.

